# G-quadruplexes in the RSV genome: potential anti-viral targets

**DOI:** 10.1101/2024.10.24.620035

**Authors:** Debopriya Bose, Suman Panda, Nilanjan Banerjee, Subhrangsu Chatterjee

**Affiliations:** Department of Biological Sciences, Bose Institute, Unified Academic Campus EN 80, Sector V, Bidhan Nagar, Kolkata – 700091, WB, India; Laboratoire d’Optique et Biosciences (LOB), Ecole Polytechnique, CNRS, INSERM, Institut Polytechnique de Paris, 91120, Palaiseau, France; Non-coding genome group, CEITEC, Kamenice 5, 62500, Brno, Czech Republic

## Abstract

The Rous sarcoma virus (RSV) is an onco-retrovirus that infects avian species such as the chicken (*Gallus gallus*). RSV is the first oncovirus to be described and the oncogenic activity of this virus is related to the expression of a tyrosine kinase that induces carcinogenic transformation. Interestingly, we have noted that the RSV genome contains various potential G4 forming sequences. Among these, two sequences located in the GAG and POL genes, respectively, show high G4 forming potential. Additionally, the SRC oncogene also harbours a putative G4 forming sequence. In this study, we have characterised the G4 formation and topology in these three loci in the RSV-DNA. We have found that these sequences form dynamic G4 structures in physiological conditions and such dynamicity may be associated with their cellular functions. Further, we have also established that these G4s are recognized by G4 interacting small-molecule ligands and the G4-stabilizing protein nucleolin. Binding of these ligands induces structural shifts in the G4 leading to change in structure and stability. Thus, the RSV-DNA G4s may be further studied as targets to control its infection and oncogenic effects.

## Introduction

The association of viruses to malignant diseases was discovered over 100 years ago with the identification of the Rous sarcoma virus (RSV). Peyton Rous noted that cancer was transmitted between chickens via the tumor extract even after it was filtered to remove all cells. He concluded that a virus must be the agent transmitting the disease, and this virus was aptly named the Rous sarcoma virus (RSV)^[1]^. The avian retrovirus RSV, contains two copies of its positive-strand RNA genome which are capped and polyadenylated. This RNA is injected into the host cell initiating infection. The RNA is reverse-transcribed into DNA within the host cell by the reverse transcriptase enzyme. The cDNA formed further integrates into the host genome through the action of integrase. Thereafter, transcription generates the viral mRNA, which either encodes viral proteins or is used in the synthesis of new viral particles. RSV contains four genes, namely, gag, pol, env, and src. Gag codes for the capsid protein, pol codes for the reverse transcriptase, env codes for the envelope protein, and src codes for a tyrosine kinase, respectively^[2]^.

G-quadruplex (G4) structures have become pivotal in the study of nucleic acid functions. Their biological relevance in various diseases have been characterised extensively. However, G4s are not only a feature of human cells or higher organisms. Their formation has now been established in several other life forms including bacteria^[3]^ and archaea^[4]^. Their functional roles have recently been highlighted even for viruses such as HIV (human immunodeficiency virus)^[5,6]^, hepatitis C^[7]^, and the human coronavirus^[8]^. In fact, G4s in the viral genome have been investigated as potential therapeutic targets to inhibit the viral life cycle. This is especially true in the case of RNA viruses as their genome is folded into various secondary structures due to the absence of a complimentary strand^[9]^. RNA-G4s are also more stable^[10]^ than their DNA counterparts and are limited to the parallel conformation^[9]^. Only a couple exceptions to this observation have been noted^[11,12]^. RNA viruses pose a massive burden in health care due to their rapid mutation rates. G4s in the viral genome are mostly located within the 5’- and 3’-UTRs or the ORFs. Deregulated formation of these G4s may inhibit essential steps such as replication, transcription, and translation^[9]^. The virus-host relationship was recently studied in terms of their potential quadruplex forming sequences (PG4s), and it was found that viral G4s have evolved to allow optimal use of the host cellular machinery^[13]^. Additionally, the base composition of specific viral G4s may be integral in the maintenance of latent infections and may serve as scaffolds for the binding of host proteins^[13]^. The spatio-temporal formation of G4s in the viral nucleic acids is therefore a major factor in controlling the viral life cycle. This also points to the roles of host G4 helicases which must be essential to allow G4 unwinding at the proper time. Thus, the stabilization of viral G4s have been proposed as a potential antiviral therapy^[9]^.

Unlike many other viruses which affect animals, RSV is also an oncovirus. The src gene of RSV codes for a tyrosine kinase, which acts as an oncogene due to non-specific phosphorylation of targets. The src gene was probably acquired by RSV via recombination between avian viruses and the host cellular src (c-Src). The result of this recombination was the viral src (v-Src) gene. This tyrosine kinase when over-expressed, induces a variety of molecular events, thus causing oncogenic transformation^[14]^. Transformation by v-Src causes altered expression profiles of several cancer associated genes in chicken cell lines. Additionally, 42 genes correlated to poor prognosis in lung and breast cancer have been found to be regulated by v-Src induced oncogenic transformation^[15]^.

While, RSV has been analysed for over a century, the formation of G4s in the RSV genome has not yet been studied. Interestingly, we have found potential G4 forming sequences in three of the four RSV genes. The gag, pol and src genes have potential G4 forming sequences which may have important biological functions. We have studied the structural topology of the DNA forms of these G4s via several biophysical assays. The circular dichroism (CD) spectra of the G4 forming DNA fragments were further used for principal component analysis (PCA) and singular value decomposition (SVD) to obtain details of the secondary and tertiary structure of each G4^[16]^. The data revealed that the GAG-G4 formed a parallel structure, the POL-G4 was a mix of parallel and anti-parallel forms and the SRC-G4 was mostly parallel with some hybrid counterpart. The structure of the RNA G4s are almost always parallel so the RNA structures were not studied in these assays. Since, viral G4s are also potential therapeutic targets, we have studied the interaction of these G4s to small-molecule ligands. We have used two G4-specific ligands, namely, TMPyP4 (5,10,15,20-Tetrakis-(N-methyl-4-pyridyl)porphine) and Braco-19 (N,N′-(9((4(dimethylamino)phenyl)amino)acridine-3-6-diyl)bis(3-(pyrrolidine-1-yl)propenamide)). These ligands have also been investigated as probable anti-viral compounds due to their role in G4 stabilization^[9]^. The binding was tested by spectroscopic techniques and additionally the effects of the binding interaction on the G4 structure was also studied. Finally, we have studied the binding of the RSV-DNA G4s to nucleolin (NCL) via electrophoretic mobility shift assay. Nucleolin is a well-known G4 stabilizing protein that has been implicated in the pathogenesis of other viruses^[17]^. In a nutshell, our studies clearly indicate the presence of stable G4 structures in the RSV genome and further the interaction of these G4s to small-molecule ligands and proteins lead to structural alteration. Thus, these G4s may be potential targets in controlling RSV infection and oncogenicity.

## Materials and Methods

### Oligonucleotide sequences

All DNA sequences used in this study were procured from Integrated DNA Technologies (IDT). The sequences were obtained as lyophilised stocks. For spectroscopic experiments, DNA samples were dissolved in water to obtain 1mM stock solutions. The fluorescently labelled samples for DNA footprinting analysis were dissolved in water to obtain 100μM stock concentration. The samples used in this study are tabulated as follows:

### Bioinformatic analysis

The RSV genome sequence of the Prague C strain was obtained from the NIH database (GenBank: J02342.1). The genome sequence and annotation was reported by Haseltine *et al*^[18]^. The sequence obtained was then analysed using the G4 predicting softwares, QGRS mapper^[19]^ and G4 Hunter^[20,21]^.

### CD-spectroscopy

The CD spectra of GG4, PG4, and SG4 were recorded using the Jasco-J815 CD spectrophotometer (Jasco International Co. Ltd). 20μM of each DNA was dissolved 10mM sodium-phosphate buffer at pH 7 and 100mM KCl. The samples were then annealed by heating at 95°C for 5 minutes, followed by gradual cooling to room temperature. The CD spectra was recorded in the 220nm-330nm range. The data was recorded at 1nm intervals and after baseline correction. The scan speed was set at 100nm/min.

### UV-spectroscopy and melting

G4 formation by the sequences under study were further confirmed by studying the absorbance of each sequence. For this purpose 2μM of each DNA was dissolved in 10mM sodium-phosphate buffer (pH 7) and 100mM KCl. The samples were then annealed as above. Further, UV-melting was performed using the Cary 60 UV-Vis spectrophotometer by Agilent. The absorbance of each sample was recorded in the 230nm-300nm range at 20°C-90°C. The data was recorded at 5°C intervals. The absorbance at 295nm with increasing temperature was then plotted as this is indicative of G4 formation^[22,23]^. The absorbance at 295nm was also used as a measure of thermal stability and was used to calculate the Tm value. The following calculations were made to determine the Tm value The fraction of folded at any temperature, T =(absF – absT)/(absF -absU),

Where,

absF is the absorbance of the fully folded state (at 20°C),

absU is the absorbance of the fully unfolded state (at 70°C, above which absorbance begins to increase),

absT is the absorbance at temperature T.

The fraction of folded so obtained is plotted against temperature and the graph obtained is fitted using the Sigma Plot software to obtain the Tm value.

### PCA and SVD analysis of CD data

The PCA (principle component analysis) and SVD (singular value decomposition) analysis of the CD spectral data was performed in accordance with the Del Villar-Guerra et al^[16]^. For this purpose, the CD data was first normalized to Δε (M^-1^·cm^-1^) =θ/(32980*c*l), where, Δε is the molar circular dichroism, θ is the ellipticity in millidegrees, c is DNA concentration in mol/L, and l is the path length in cm. The molar CD spectra so obtained was analysed against a reference library of 23 G4s of known conformation. The G4 structures were grouped by PCA followed by HCPC (hierarchial clustering of principal components). This was performed using the R software and the conformation of the unknown sequences was analysed from this data. The secondary and tertiary sequences were further analysed by SVD analysis by comparing the spectra against the reference library.

### DMS protection assay

To further identify the residues involved in G4 formation, we performed the DMS protection assay. Briefly, DMS methylates the N7-residues of guanine residues and the DNA strand is further cleaved at the methylated positions by piperidine. However, if the guanine residue is involved in G4 tetrad formation, the N7 residues form Hoogsteen hydrogen bonds with the N2-H moiety. This protects the N7 residue from DMS induced methylation and further cleavage. The participation of guanine residues in tetrad formation can then be analysed by gel electrophoresis of the DNA sequence in conditions allowing and inhibiting G4 formation^[24]^. The DMS protection assay was performed using 70nt sequences containing the G4 loci of interest. 10μM of each DNA sequence was then annealed in water or sodium-phosphate buffer along with KCl. Thereafter, the annealed sequences were used for the DMS reaction. Each reaction contained 500nM DNA in the presence or absence of KCl, 20μl of 1% DMS, and water (upto 100μl). The sample was then incubated for 2 minutes exactly at room temperature after which the reaction was stopped using a mixture of β-mercaptoethanol and 3M sodium acetate (pH 7). The DNA was then precipitated and washed with ethanol followed by immediate lyophilisation. Further, the modified DNA was cleaved by heating with 1M piperidine for 20 minutes followed by immediate lyophilisation. The DNA was then washed thrice with water. Additionally, an A/G ladder was developed by treating the DNA with 4% formic acid and incubating at 37°C for 30 minutes followed by immediate lyophilisation. The A/G ladder sample was also treated with piperidine and processed as above. All samples were finally dissolved in formamide dye and heated for 10 minutes at 95°C followed by snap-chilling. The samples were then electrophoresed in 15% PAGE gel, containing 8M urea. The gel was finally imaged using the Typhoon Trio+ phosphorimager from GE Healthcare. The gel image was analysed with the Image J software to unequivocally identify the intensity of each band.

### Fluorescence spectroscopy and titration

We have also assayed the binding of the G4 sequences under study to the G4 specific small-molecule ligands, TMPyP4 and Braco-19. For this purpose, we have used fluorescence spectroscopy. For this purpose, we have used the Hitachi spectrophotometer, F-7000 FL. 5μM TMPyP4 or 5μM Braco-19 were titrated with increasing concentrations of the G4 DNA under study. The DNA was annealed in sodium-phosphate buffer (pH 7) in the presence of KCl prior to the titration experiment. TMPyP4 was excited at 435nm and emission spectra was recorded in the 600nm-750nm range. Braco-19 was excited at 300nm and emission spectra was recorded at 350nm-550nm. The data was recorded using a cuvette of 1cm path-length in a right-angle geometry. Data was recorded at 1nm intervals and scan speed was set at 100nm/minute. All spectra were corrected for dilution before analysis.

### CD-spectroscopic titration

The binding of the G4 forming DNAs under study to Braco-19 was also assayed by CD spectroscopy. 20μM of the DNA sequences were annealed in the presence of 10mM sodium-phosphate buffer (pH 7) containing 100mM KCl. The DNA samples were then titrated with increasing concentration of Braco-19 to study whether ligand binding affects G4 structure. The Jasco-J815 CD spectrophotometer (Jasco International Co. Ltd) was used for this study. The CD spectra were recorded in the 210nm-310nm range and data was collected at 1nm intervals post baseline correction. The scan speed was set to 100nm/minute.

### Homology modelling

The similarity between the chicken and human nucleolin proteins was checked by homology modelling via SWISS-MODEL^[25–29]^ and I-TASSER^[30–32]^ softwares. Since only the structure of the RNA-binding domains (RBDs) 1 and 2 of human nucleolin are available (PDB ID: 2KRR), the homology modelling was performed only for these domains. 2KRR was used as a template for modelling in both servers and the models obtained were aligned using the Pymol software.

### Electrophoretic mobility shift assay (EMSA)

Binding assay was performed with the G4 forming sequences and purified human nucleolin. Nucleolin was procured from Biobharati Pvt. Ltd., India. EMSA was performed using a previously reported protocol^[17]^ with slight modifications. Briefly, 10μM DNA samples were annealed in a buffer containing 10mM Na-cacodylate buffer (pH 7) in the presence of 100mM KCl and 15% PEG 300. Thereafter, the binding buffer for EMSA was prepared by combining 20mM Tris-Cl, 30mM KCl, 1.5nM MgCl_2_, 1mM DTT, 8% glycerol, 1% protease inhibitor cocktail, and 15% PEG 300. 500nM of DNA was incubated in this buffer at 37°C for 30 minutes in the presence or absence of 3.75μg of human nucleolin. The samples were then electrophoresed in a 6% non-denaturing PAGE at 40V. Finally, the gel was photographed using the Typhoon Trio+ phosphorimager from GE Healthcare.

### Statistical analysis

All experiments were performed thrice to ensure that the results were significant. Average values and standard errors (SE) of the data were calculated using Microsoft Excel. The graphs were plotted as average ± SE using the Sigma Plot software.

## Results and discussion

### G-quadruplexes are formed in RSV DNA

The RSV genome consists of its positive strand RNA that is reverse-transcribed to form the DNA sequence which integrates into the host genome and is further transcribed to form the mRNA. The genome sequence of the Prague-C strain of RSV was first analysed using the softwares QGRS mapper and G4 hunter. The RSV genome has a high proportion of potential G4 forming sequences. QGRS mapper predicted 80 individual G4 forming sequences while G4-hunter predicted 43. The highest scoring G4 forming sequences among these were located within the GAG and POL genomic regions. The predicted G4 forming sequences in these genes are tabulated in Table 3. Further, we analysed the G4s in the SRC gene sequence. While this sequence did not show any G4 with high propensity, a sequence with low G-score was present. Due to the importance of this gene in promoting carcinogenic events, we chose to study this sequence. Impeding transcription or translation of the SRC G4 may be instrumental in curbing the oncogenicity of v-SRC.

**Table 1:**
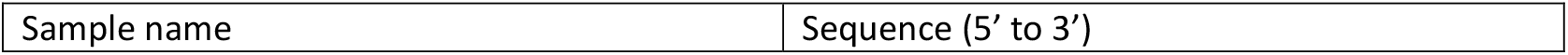
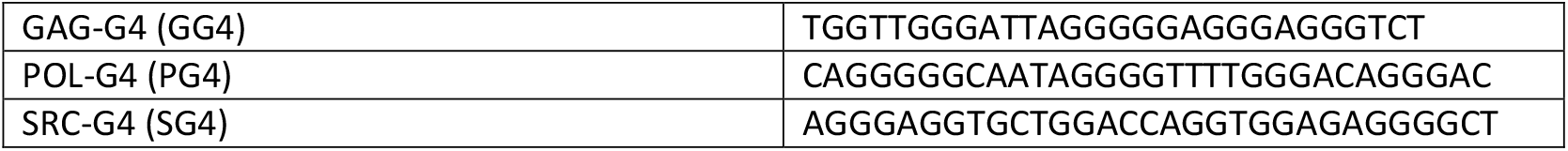
DNA sequences for spectroscopic analysis.

**Table 2:**
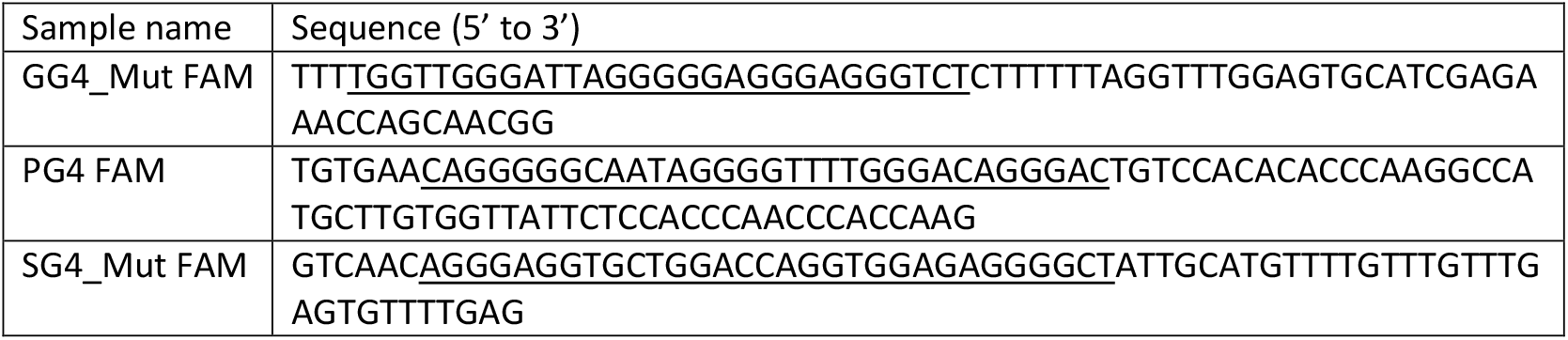
Sequences for DNA footprinting and EMSA (Tagged with 6-FAM at 5’-end)

**Table 3:**
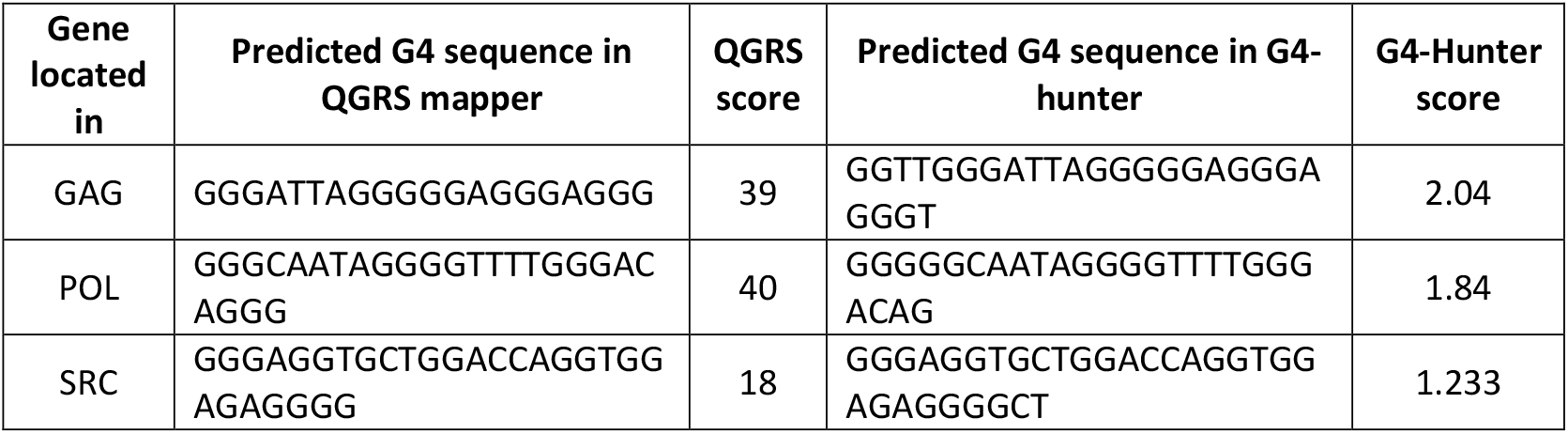
Potential G4 forming sequences in the RSV genome and their G-scores.

Based on these results, we proceeded to analyse the G4 formation by the predicted sequences experimentally (Sequences chosen for experiments can be found in the materials and methods section). For this purpose, we chose to perform CD- and UV-spectroscopy. CD-spectroscopy provides a method to identify secondary structures in DNA via distinctive spectra obtained for various structural forms. In the case of G4s, various CD spectra may be possible depending on the strand orientation within the G4. Parallel structures are generally characterized by a positive ellipticity peak close to 260nm and a negative peak close to 240nm. On the other hand, anti-parallel G4s shows a positive peak around 295nm and a negative peak around 260nm. Finally, hybrid G4s show two positive peaks close to 260nm and 290nm along with a single negative peak at 240nm^[33,34]^. Upon analysis of the CD-spectra of GG4, PG4, and SG4 it was found that GG4 forms a parallel-G4 structure. However, the spectra obtained for PG4 and SG4 was more complex. Both sequences show a positive peak around 260nm, however, the peaks are quite broad and it was difficult to predict their topology unequivocally. Therefore, we performed UV-spectroscopy to confirm the formation of G4 structures in the sequences under study. UV-melting experiments are another way of confirming the formation of G4s in a sequence of interest. While UV-melting of duplex or other conformational forms of DNA leads to a hyperchromic transition with increasing temperature, G4s show a distinctive trend. All G4 forming sequences show a hypochromic transition of absorbance with increasing temperature at 295nm^[22,35]^. As expected, all sequences showed a hypochromic transition at 295nm confirming G4 formation. The UV-melting data was also used to calculate the stabilities of the G4 structures. The fraction of folded at each temperature was calculated (see materials and methods section) and plotted against temperature. The graph showed the best-fit to the sigmoidal equation and the average temperature was obtained from the Sigma Plot software. All sequences show a Tm above physiological temperatures of the RSV host, suggesting the possibility of formation of these structures *in vivo*. The formation of G4 structures by these sequences have also been confirmed via NMR-spectroscopy. G4 structures are characterized by the formation of peaks between the 10-12.5ppm regions in the NMR spectra. All sequences under study show the formation of these peaks, as expected (Supplementary information, Fig 1).

**Figure 1:**
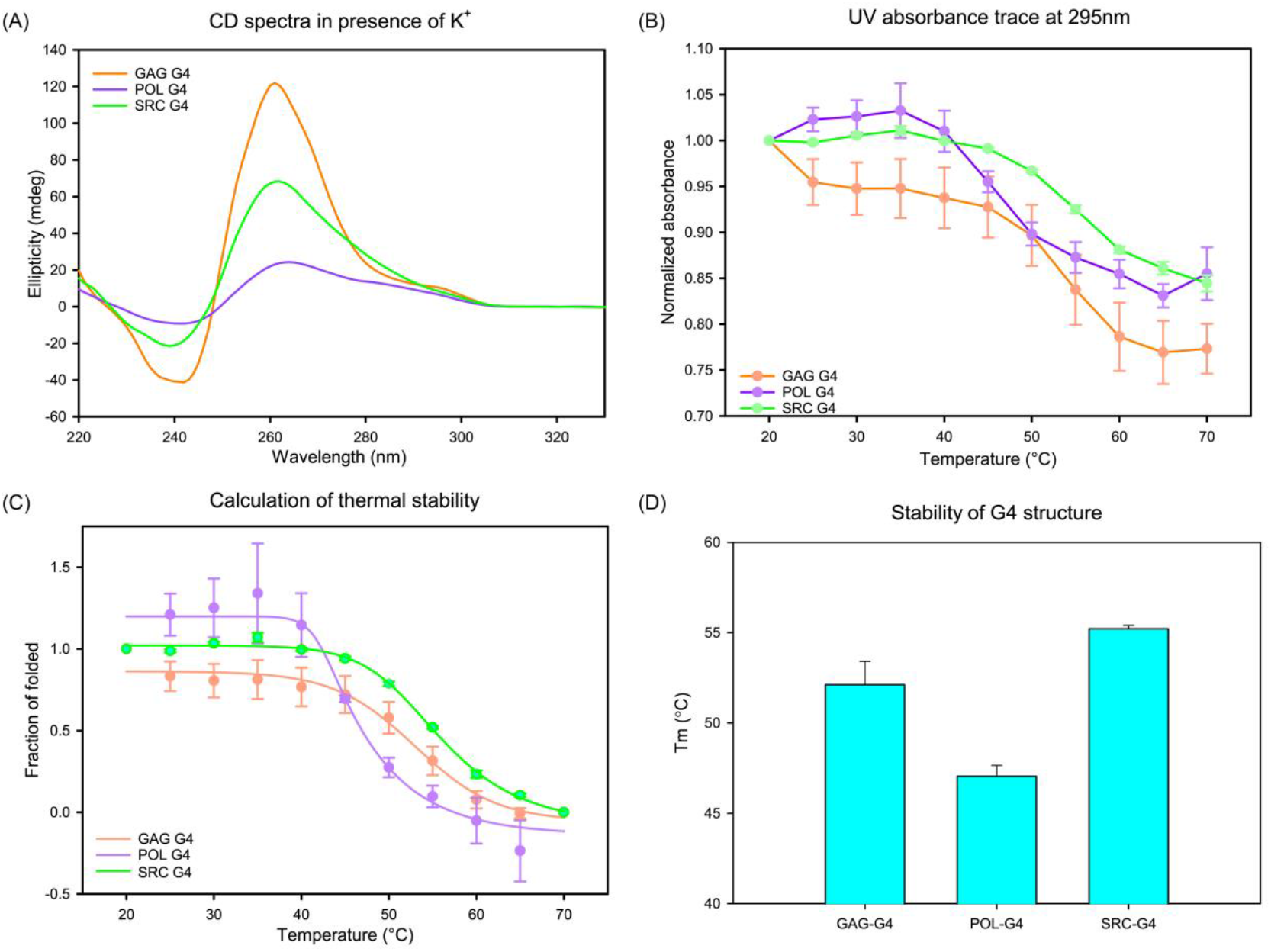
Formation of G4 structures by sequences in the RSV-DNA. (A) CD-spectra of GG4, PG4, and SG4 show the formation of parallel G4 by GG4, but the structures of PG4 and SG4 are ambiguous. (B) Confirmation of G4 formation by the sequences under study by UV-melting assay. All sequences show a hypochromic transition of absorbance with increasing temperature at 295nm. (C) Calculation of the fraction of folded at each temperature from UV-melting data and plotting against temperature. The curves showed the best-fit to sigmoidal nature and was used to calculate the average melting temperature. (D) The Tm obtained from curve-fitting was plotted for each G4. All sequences are stable above physiological temperatures of adult chickens, suggesting that these G4s may form *in vivo*.

### Confirming the structural topology of the RSV-DNA G4s

While G4 formation was confirmed from the above experiments, the structural topology of SG4 and PG4 remained ambiguous. So, we sought to better understand the structures using the CD-data obtained. It is well-known that the CD-spectra of proteins can be analysed to obtain the percentages of each secondary structural form in the protein. A similar method has now been developed to characterize the structural forms of G4 structures^[16]^. This method involves analysis of the normalized CD-spectra (Supplementary information, Fig 2) against a reference library of confirmed G4 structures. The reference library for this assay has been developed by Del Villar-Guerra et al. using 23 solved G4 structures. Briefly, the authors analysed the CD spectra of these 23 G4s in conditions ideal to which the structures were solved in and clustered the data via principle-component analysis using the R software. This generated three distinct populations of G4s depending on structural forms. Further, the data obtained were also subjected to singular value decomposition (SVD) and least-square fitting via R software to obtain a quantitative structural analysis. The structural features of GG4, PG4, and SG4 were analysed using a Shinny web R program provided by the original authors which performs a detailed analysis of the structural properties of each G4. The analysis shows that the GAG-G4 is completely of a parallel nature as predicted, while POL-G4 is a mix of the parallel and anti-parallel forms. Finally, the SRC-G4 is dominantly parallel in nature with some amounts of hybrid and anti-parallel structural variants. The details of the secondary and tertiary structural parameters of these G4s can be found in the Supplementary information, Fig 3.

**Figure 2:**
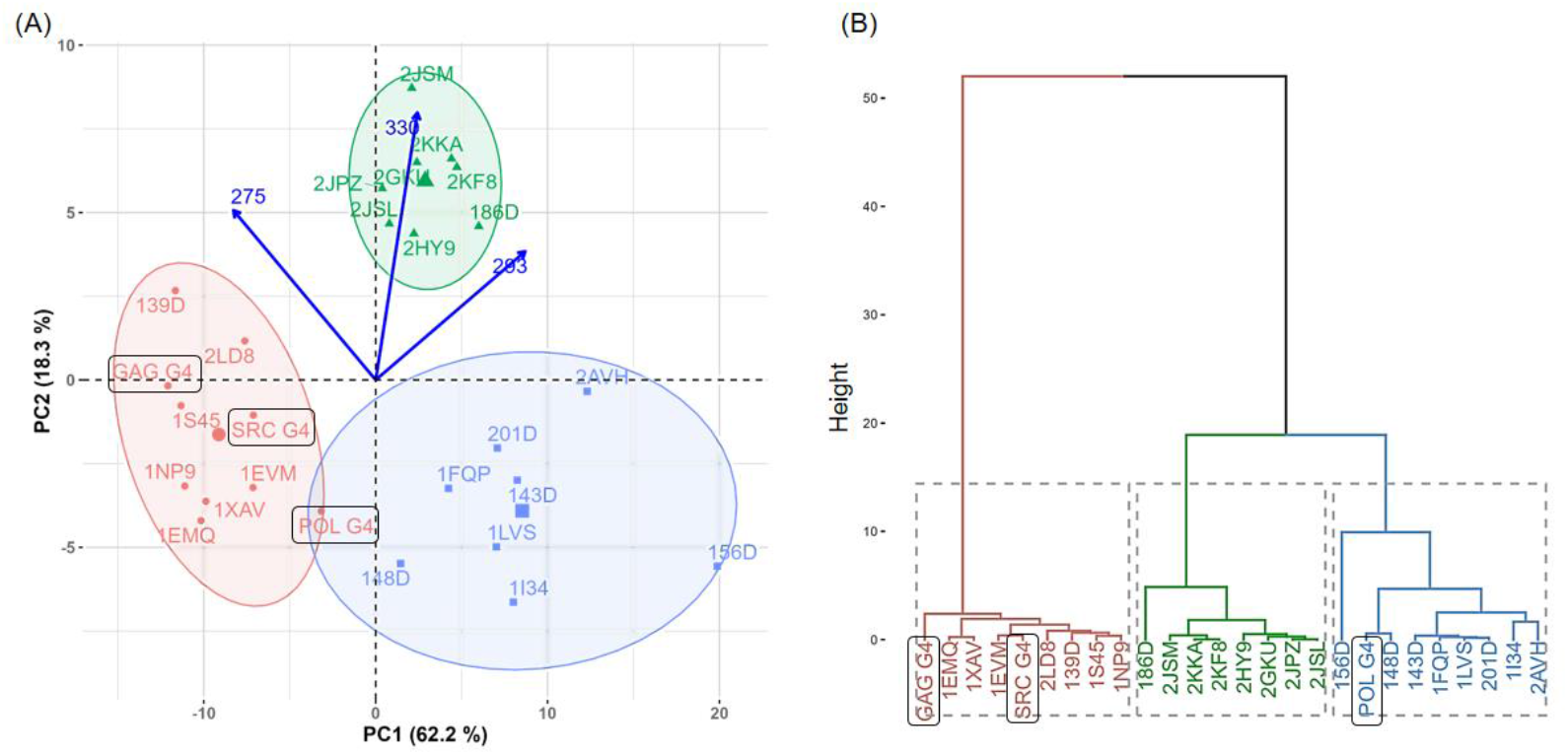
Structural analysis of GG4, PG4, and SG4 via analysis of CD-spectra. (A) PCA of GG4, PG4, and SG4 with the reference CD spectra. The PDB IDs of the reference spectra are present in the plot, showing the clustering of GG4, PG4, and SG4. The red cluster represents parallel structures, the blue cluster represents anti-parallel structures and the green cluster represents hybrid structures. (B) Hierarchical clustering on principal components (HCPC) shows the dominant structural parameters of each G4. The colour coding is same as that of the PCA analysis.

### Prediction of the G4 structures in the RSV-DNA

Once the structural topology of the G4s under study were unequivocally assigned via computational analysis, we proceeded to characterize the structures of the G4s further. For this purpose, we performed DMS footprinting using DNA sequences tagged with the fluorophore FAM at the 5’-end. Formation of a G4 structure requires the stacking of multiple G-tetrads. Each G-tetrad is formed by four guanine residues bonded to each other via Hoogsteen hydrogen bonds forming a square planar arrangement. In a G-tetrad, each guanine residue acts both as a hydrogen bond donor and acceptor and forms the N1-H----O6 and N2-H N7 bonds. DMS (dimethyl sulphate) methylates the N7 residues of guanine when free but is unable to methylate the N7 residues involved in Hoogsteen hydrogen bonds. Thus, G4 formation protects DNA from DMS induced methylation at guanines involved in tetrad formation. Further, these sites are also protected from piperidine induced cleavage^[36]^. Thus, guanines participating in G4 formation can be identified from DMS footprinting.

Initially, we performed DMS footprinting experiments using 70nt sequences containing the GG4, PG4, and SG4 loci. The flanking sequences in these nucleotides were same as in the wild-type RSV sequence. However, it was observed that only PG4 showed KCl induced protection from cleavage in these conditions while GG4 and SG4 showed no protection (Supplementary information, Fig 4). This is in contrast to the data obtained thus far confirming G4 formation in the sequences under study. Therefore, we analysed the sequences used and observed that both the GG4 and SG4 loci are followed by C-rich regions which might form hairpin structures thereby hampering G4 formation. Such G4-hairpin switches may also be important in regulation of the expression of these genes. Interestingly, the SG4 sequence is followed by a potential i-motif forming loci which may also be crucial in SRC gene transcription. Sequences showing such structural diversity and switching properties have been found to be important in various genomic loci and need to be studied further^[37–39]^. To confirm this, we mutated the flanking sequences of GG4 and SG4 to replace the cytosines with thymine residues (Supplementary information, Fig 4). DMS footprinting with these sequences revealed KCl-induced protection in all sequences as expected. The data revealed that seven guanine residues in the GG4 sequence participate in tetrad formation. This suggests that the GG4 structure *in vitro* is intermolecular in nature as the formation of a unimolecular parallel G4 would require the protection of at least eight guanines. The GG4 structure was deciphered to be parallel from the CD-data and this information has been combined to predict a probable model of the GG4 intermolecular structure. On the other hand, PG4 and SG4 show the protection of twelve and fourteen guanines, respectively. The PG4 structure is a mix between parallel and anti-parallel forms and therefore two structures have been predicted. The PG4 structure is quite unique as the protected sequence contains three G-runs instead of four. This introduces certain constrictions in the formation of the G4 structure as only two loop forming sequences are available. Considering these conditions, we predicted that the PG4 structure forms a (4n-1) G4 with a G-vacancy in one tetrad. However, it is possible that this vacancy is fulfilled by flanking adenine residues stabilizing the structure. Similar, adenine-driven G4 structures have also been found elsewhere^[40]^. It is also possible that some loops in this structure are devoid of nucleotides. It is probable that the POL-G4 structure has many alternative structures. In the model shown in figure 3, we have included G11 in the tetrad plate. However, it is equally probable that G11 may be present in the first loop thus allowing G14 to participate in tetrad formation. Finally, the SG4 structure is also quite unique in its guanine distribution. As expected, the SG4 structure is probably extremely dynamic in nature as suggested by the protection of fourteen guanine residues. We have predicted two structural alternatives of the SG4 parallel structure, however, it is quite probable that many other structural alternatives are present, which may be parallel, hybrid or anti-parallel in nature. The G25 residue is equally probable to be present in tetrad formation as G23 and this itself would give two more structurally equivalent G4 forms. Thus, the G4s in RSV-DNA are potential sites for molecular switching to various structural forms suggesting their potential importance in regulation of transcription and translation of these genes.

**Figure 3:**
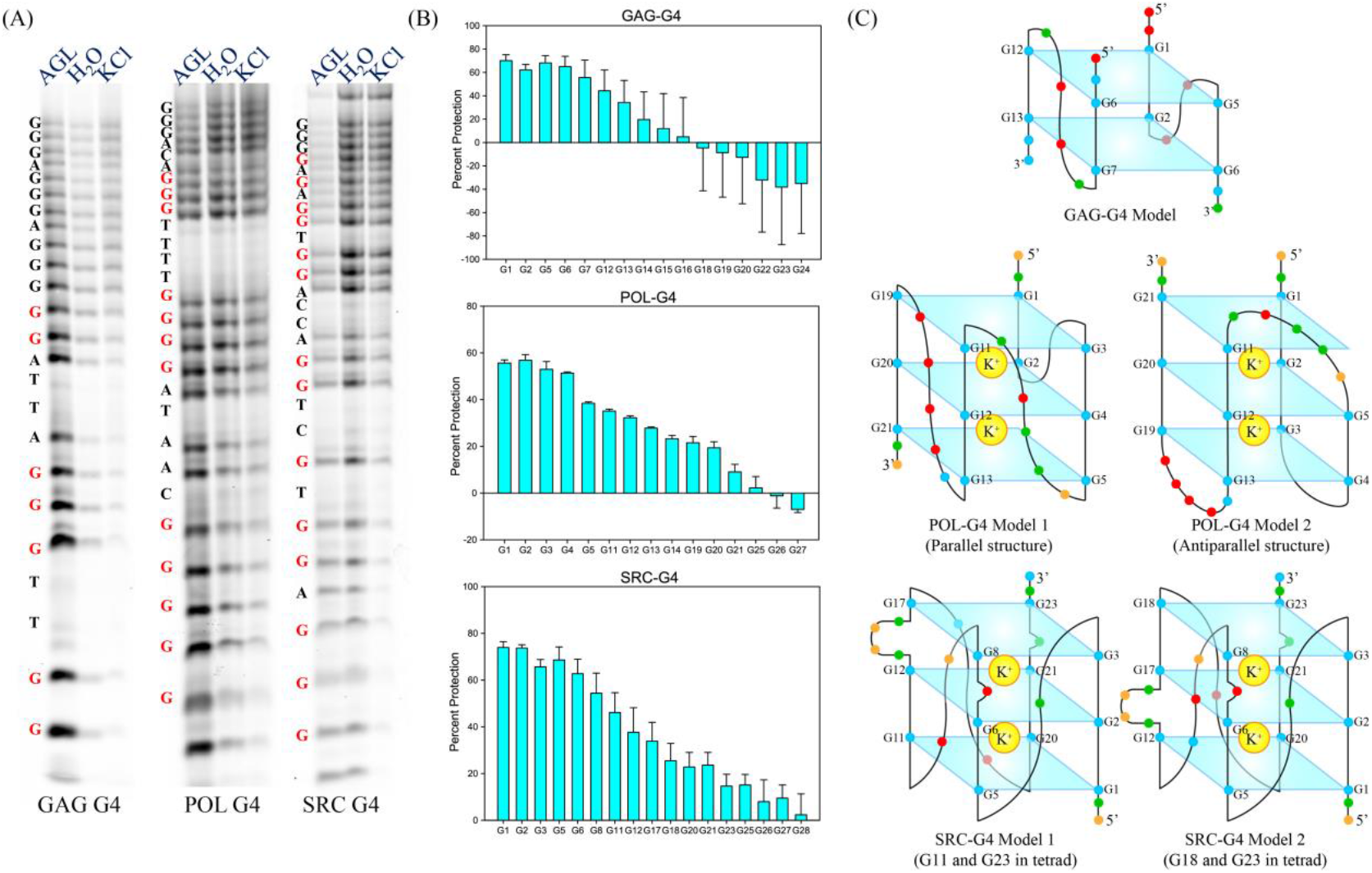
DMS-protection assay of RSV-DNA G4s. (A) Gel electrophoresis showing the protection of guanine residues in the presence of KCl suggesting their involvement in tetrad formation. (B) Calculation of percentage of protection of each guanine residue from gel electrophoresis images to identify the guanines involved in tetrad formation. (C) Probable models of the G4 structure predicted from DMS-footprinting data. Blue dots in the models show guanine residues, green dots show adenine residues, red dots show thymine residues and yellow dots show cytosine residues, respectively.

### Binding of G4 structures in the RSV-DNA to G4-specific small molecule ligands

Targeting G4 structures within cells requires their stabilization or destabilization via external agents such as small-molecule ligands or aptamers. The formation of G4s have now been predicted in all human viruses and further confirmed in several viruses^[41]^ such as HIV (human immunodeficiency virus)^[42,43]^, corona virus (SARS-CoV-2)^[44]^, and Zika virus^[45]^. As previously mentioned, these G4s play important roles in hijacking the host apparatus by acting as sites for protein binding. However, spatio-temporal unwinding of these structures are necessary for completion of essential processes such as transcription and translation. The G4s in viral genomes thus act as potential targets for inhibition of the viral life-cycle^[46]^. Various G4 specific small-molecule ligands such as TMPyP4 and Braco-19 have therefore been investigated as anti-viral therapeutics^[47]^. While these compounds bind to cellular G4s as well, the anti-viral activity stems from an abundance of viral nucleic acids within infected cells.

We have thus investigated whether the G4s in the RSV-DNA can be recognized by such G4 specific ligands. For this purpose, we have performed fluorescence- and UV-spectroscopic titration experiments using the sequences under study. Binding of TMPyP4 or Braco-19 to GG4, PG4, and SG4 were first studied by fluorescence spectroscopy wherein the ligands were titrated with increasing concentrations of DNA annealed in buffer in the presence of KCl. In the case of TMPyP4, we have observed that increase in GG4 concentration leads to a hypochromic transition followed by peak-shift (red-shift) and hyperchromicity at higher DNA concentrations. In the case of PG4, TMPyP4 fluorescence increased upon titration (hyperchromic shift) along with a slight red-shift. Finally, SG4 led to hyperchromicity at low concentrations followed by hypochromic shift and red-shift as well. In the case of Braco-19, hypochromic shifts were observed upon titration with each DNA sequence followed by hyperchromicity in the case of PG4 and SG4. This is also associated with alteration in the peak nature and peak-shifts. Binding of these compounds to the G4s was also investigated via UV-spectroscopic titrations in a similar experiment. Peak shifts suggesting interaction was also observed in this experiment (Supplementary information, Fig 5) and the Kd for the interaction between Braco-19 and RSV-DNA G4s was also calculated from this data (Supplementary information, Fig 6). Thus, both TMPyP4 and Braco-19 interact with the G4s under study.

**Figure 4:**
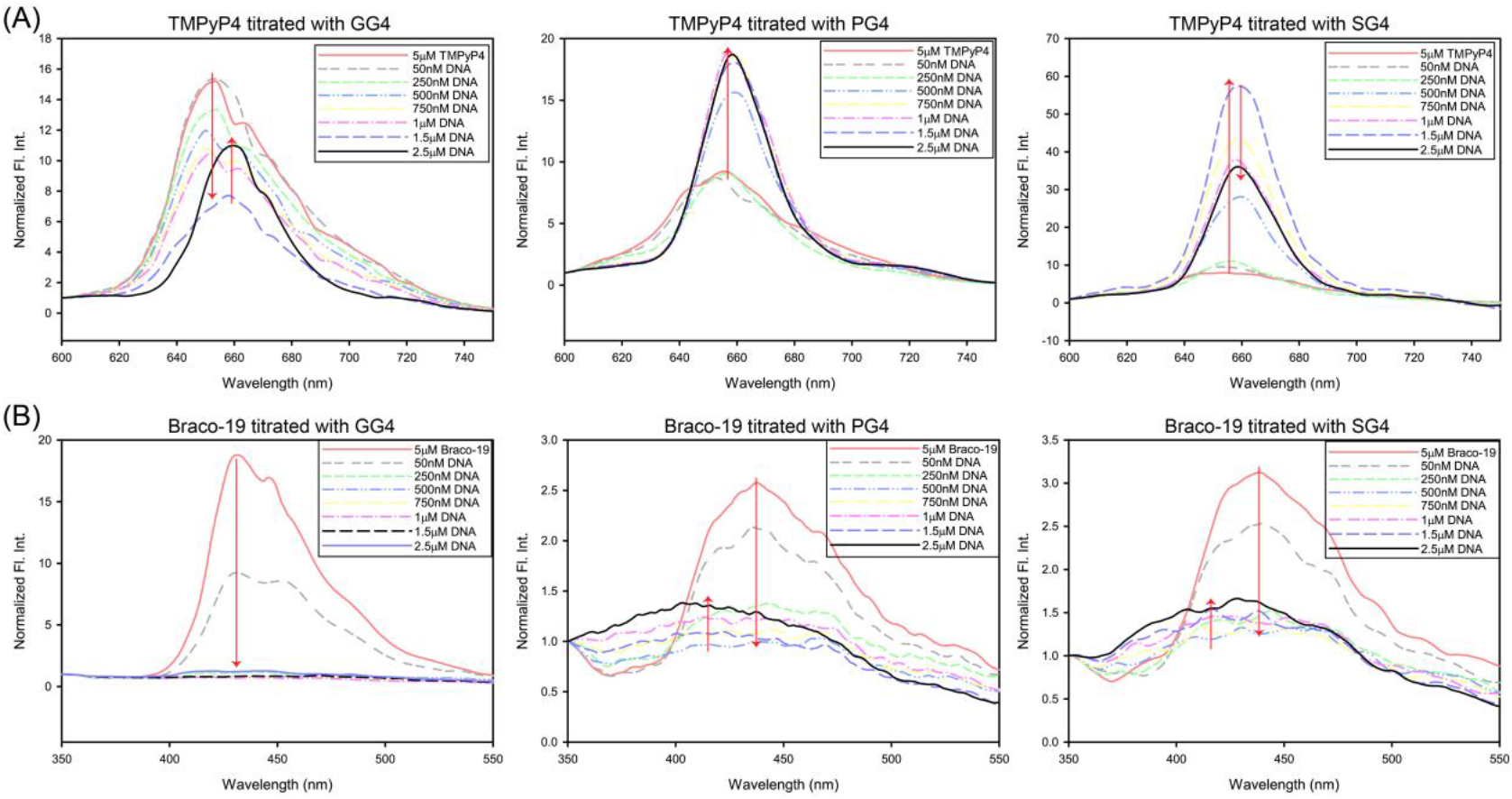
Fluorescence spectra showing the alteration of peaks for (A) TMPyP4 or (B) Braco-19 upon titration with the G4 DNA sequences under study. Titration with DNA leads to significant peak shifts for each ligand, suggesting interaction with the G4 structures. Structural differences of the individual G4s leads to different effects on ligand spectra, as expected.

### Effect of ligand binding on G4 structures in the RSV-DNA

Once the binding of TMPyP4 and Braco-19 was confirmed to the G4 structures under study, we proceeded to characterize the effect of this binding on the G4 structures. For this purpose, we have used CD and NMR (nuclear magnetic resonance) spectroscopy. As previously mentioned, CD spectroscopy is an important method to understand the secondary structures of biological macromolecules including DNA. For this analysis, increasing concentrations of Braco-19 were added to the G4 sequences annealed in the presence KCl in Na-P buffer. As expected, this led to shifts in the CD-spectral peaks of the G4 structures. In the case of PG4 and SG4, it was observed that increasing ligand concentration led to decrease in the peak at 260nm and increase in the peak at 295nm. Additionally, for PG4 ellipticity increases at 240nm upon ligand titration. This suggests that the PG4 conformational equilibrium shifts more toward the anti-parallel conformation upon binding to Braco-19. However, in the case of SG4, the ellipticity at 240nm also decreases further suggesting that in this case the conformation shifts towards the hybrid topology. In the case of GG4, no overall conformational shift is noted. However, the parallel G4 may be destabilised leading to decrease in ellipticity at 260nm and slight increase in ellipticity at 240nm. Thus, ligand binding causes structural changes in all the G4s being studied and this may interfere with the roles of these G4s in the progression of RSV infection. The structural alteration of G4s via Braco-19 interaction has also been confirmed by NMR-spectroscopic titration. Herein, we observed that ligand binding led to various alterations in the DNA spectra. In the case of GG4, Braco-19 leads to removal of certain peaks suggesting either destabilization or increased dynamicity. However, titration of PG4 or SG4 with Braco-19 led to the development of additional imino proton peaks in the 10-12.5ppm region corresponding to G4 formation. This suggests that these two G4s are stabilized by ligand interaction in addition to undergoing structural alterations. The NMR spectroscopy data are added in the Supplementary information, Fig 7.

**Figure 5:**
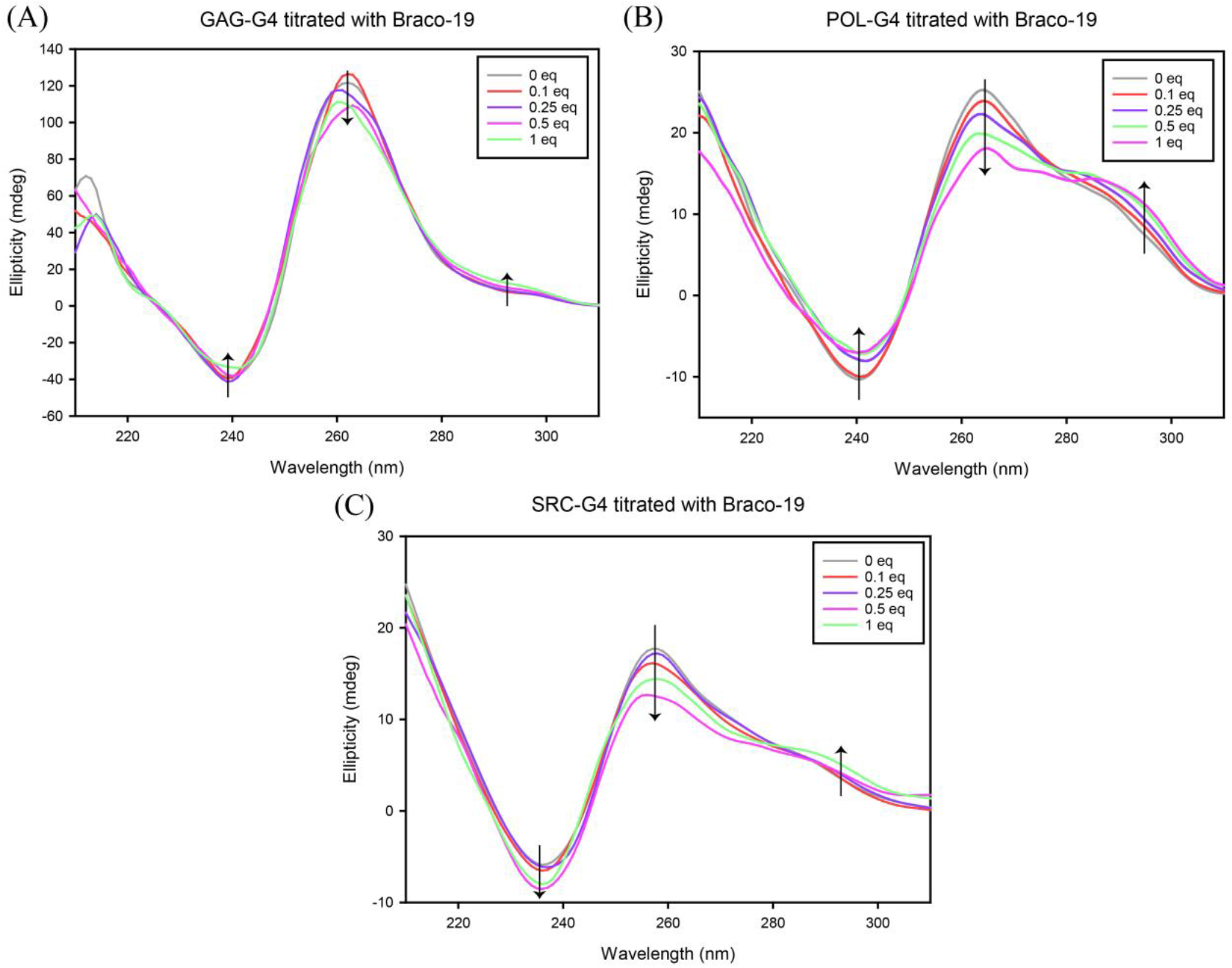
Structural changes induced by Braco-19 on RSV-DNA G4s. (A) The GG4 spectra shows decrease at 260nm and increase at 240nm as an effect of Braco-19 interaction, suggesting destabilisation of the parallel G4 structure. (B) PG4 undergoes a transition leading to increase in its anti-parallel conformation upon Braco-19 binding. The anti-parallel G4 structure shows a positive peak at 295nm and negative peak at 260nm, along with a positive peak at 240nm. Braco-19 leads to increase in the 295nm region and 240nm region while decreasing the 260nm peak, thereby promoting the anti-parallel form of PG4. (C) The SG4 spectra shows an increase in the 290nm region and a decrease in the 260nm and 240nm region as a result of Braco-19 binding. These shifts suggests that the structure transitions more toward the hybrid conformation (positive peaks at 260nm and 290nm, negative peak at 240nm) due to interaction with Braco-19.

### Nucleolin binds the RSV-DNA G4s which may lead to G4 stabilization in vivo

Nucleolin is a well-studied G4 binding protein which stabilises G4 structures and plays important roles in nucleic acid chaperoning^[48]^. In addition to the role of this protein in the maintenance of vital cellular activities such as transcription, this protein has also been reported to be involved in the stabilization of G4s in the HIV-1 LTR promoter which leads to silencing of viral transcription^[17]^. The role of nucleolin has also been demonstrated in the regulation of various viral life cycles^[48]^. Therefore, we hypothesized that nucleolin may be one of the host proteins involved in G4 mediated regulation of RSV infection. To corroborate this hypothesis, we first checked the structural similarity of the human and chicken nucleolin. For this purpose, we used the reported structure of the human nucleolin (RBD 1 and 2, PDB id: 2KRR) to model the probable structure of the chicken nucleolin. Homology modelling was performed via two softwares, SWISS-MODEL and I-TASSER, and high degree of alignment was observed in both cases. Thereafter, performed EMSA experiments to check the interaction between the G4-DNA and human nucleolin. It was clearly observed that addition of nucleolin leads to upward shift of the DNA band in all three cases confirming interaction of nucleolin with these sequences. Thus, it is probable that the G4s in the RSV-DNA play significant biological roles via its ability to interact with nucleolin (and possibly many other G4 binding proteins) in physiological conditions.

**Figure 6:**
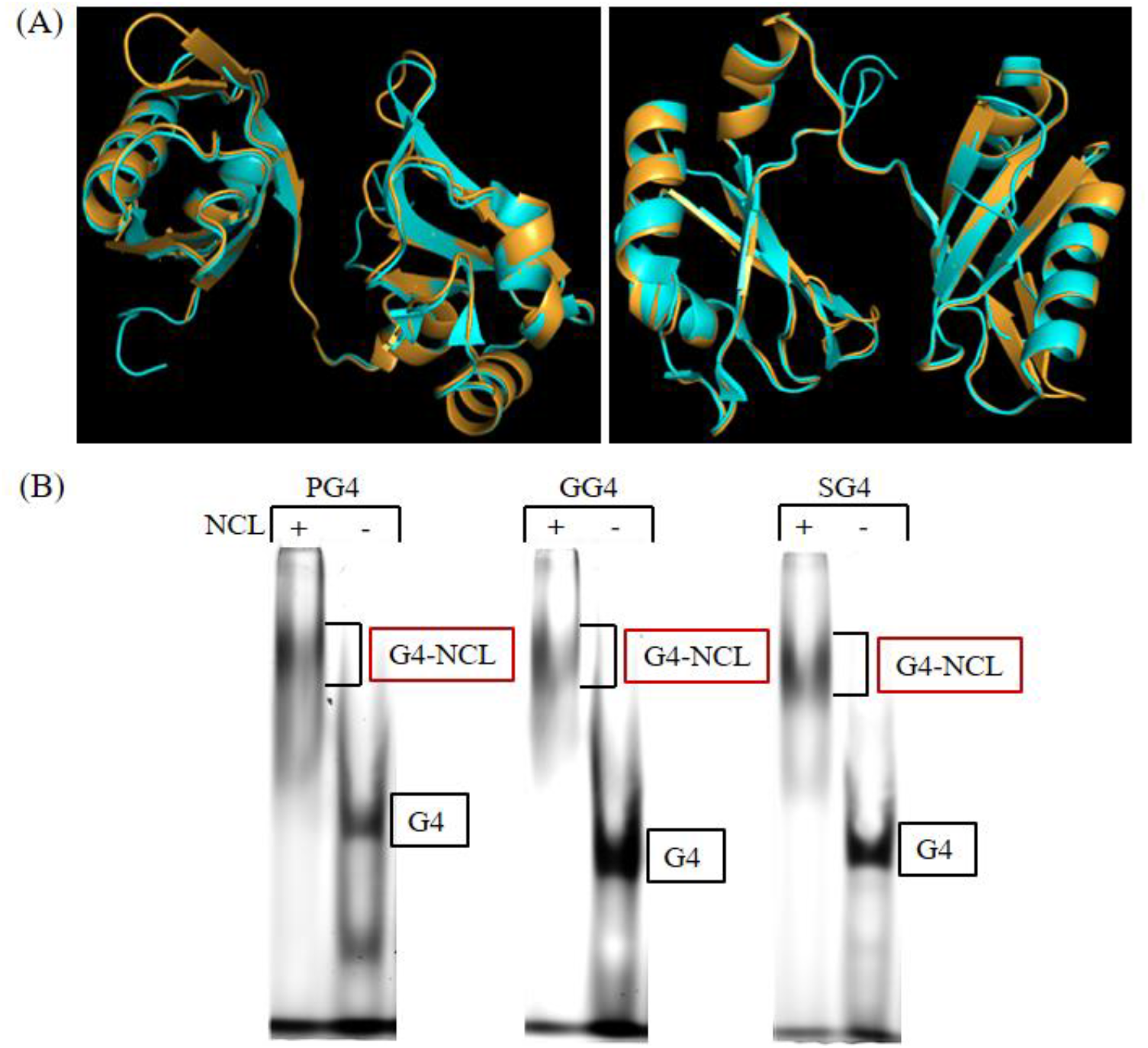
RSV-DNA G4s interact with nucleolin (NCL) in physiological conditions. (A) Homology modelling of chicken NCL (RBD 1 and 2) with respect to human nucleolin via SWISS-MODEL, shows high degree of structural similarity. The homology was also confirmed via I-TASSER and the data is available in the supplementary information, figure 8. The chicken nucleolin is shown as orange and the human nucleolin is blue. (B) EMSA shows interaction of RSV-DNA G4s with nucleolin. The G4-NCL complexes in all three cases are shifted upwards compared to the DNA only column.

### Summary

RSV is an important pathogen that causes oncogenic transformation in its host via the action of a protein kinase that it expresses. The RSV genome is reverse-transcribed into its complementary DNA, which then integrates into the host genome. This DNA thereafter serves as a template for transcription to manufacture viral proteins. The viral life-cycle can therefore be inhibited if the functional elements of this DNA is altered. In this aspect, G4s may play an important role due to their involvement in hijacking of the host machinery. Interestingly, the RSV-DNA contains multiple probable G4 forming elements among which the sequences with highest G4 forming propensity are located within the GAG and POL genes. Additionally, a sequence within the SRC oncogene also has G4 forming potential. In this study, we have verified the G4 formation by these sequences via various biophysical assays. Further, the structural topology of these G4s have also been studied by computational and biophysical methods. We have established, that GG4 forms a parallel G4 structure while PG4 and SG4 form highly dynamic G4s switching between various structural forms. Such molecular switching behaviour may also aid in the functional properties of these G4s *in vivo*, however, further studies are required to elucidate the functional properties of these elements. We have also analysed the binding of these G4s to specific small-molecule ligands and also analysed the structural changes induced by the binding of Braco-19 on the G4. Finally, we have observed that the G4 forming sequences in the RSV-DNA are recognised and bound by human nucleolin, which is highly similar in structure to the chicken nucleolin. This suggests that the G4s in the RSV-DNA may be implicated in various biological functions. In essence, these studies conclude that G4s are formed in the RSV-DNA at multiple locations and these G4s show molecular switching properties under physiological conditions. Further, these G4s are also bound by small-molecule ligands and proteins which induce structural changes. Thus, these G4s may be targetable sites for the control of RSV infection.

## Corresponding Author

*To whom correspondence should be addressed

Subhrangsu Chatterjee, Professor, Department of Biological Science, Bose Institute, Unified Academic Campus, N 80, Sector V, Bidhan Nagar Kolkata - 700091 WB India

## Acknowledgement

The authors are deeply grateful to Prof. Dr. Jonathan B. Chaires and Dr. Rafael del Villar Guerra for providing us the Shinny web R program containing the reference CD spectra of 23 G-quadruplexes, for the structural analysis.

