## Supplementary Information for "G-quadruplexes in the RSV genome: potential anti-viral targets"

### **Experimental methods:**

#### 1. 1-D NMR spectroscopy:

The NMR spectra of the DNA sequences under study were recorded using the Bruker ASCEND^TM^ 700MHz NMR spectrophotometer. It is known that G4 formation leads to the development of characteristic peaks in the 10.0-12.5ppm region due to the formation of characteristic Hoogsteen hydrogen bonds. For the spectroscopy, 300μM DNA was annealed in 10mM Na-P buffer in the presence of 100mM KCl. All samples also contained 10% D_2_O and TSP (3- (trimethylsilyl)-2,2′,3,3′-tetradeuteropropionic acid) which was used as an internal standard (0.0ppm). Spectra were recorded at room temperature and the number of scans was fixed at 128.

#### 2. UV-Vis titration experiments:

The binding of the RSV-DNA G4s to the G4 specific small-molecule ligands, TMPyP4 and Braco-19, were checked by UV-Vis titration experiments. The data were recorded using the Cary 60 UV-Vis spectrophotometer from Agilent. For this purpose, 2μM TMPyP4 was titrated with increasing concentrations of the G4s annealed in Na-P buffer in the presence of KCl. The absorbance of TMPyP4 was recorded in the 350nm-500nm range. In the case of Braco-19, 50μM Braco-19 was titrated with increasing concentration of the G4 DNAs as before. All data were corrected for dilution and plotted. The putative Kd for the interaction between Braco-19 and the G4s under study were also calculated, as Braco-19 was also used for further experiments. For this purpose, the absorbance maxima of Braco-19 was noted and the change in absorbance at this wavelength was plotted as a function of DNA concentration. The normalized absorbance for this plot was calculated as follows:

(Free ligand absorbance – absorbance at each point)/(Free ligand absorbance – bound ligand absorbance)

The normalized absorbance was plotted against DNA concentration and the graph so obtained was fitted using the Sigma Plot software to obtain the putative Kd.

#### 3. 1-D NMR titration:

The structural changes induced by Braco-19 were further studied using 1-D NMR titration experiments. The Bruker ASCEND^TM^ 700MHz NMR spectrophotometer was used to record the data. The number of scans was set at 128 and the temperature was set at 25°C. 300μM DNA annealed in Na-P buffer in the presence of KCl (containing 10% D_2_O and TSP) was titrated with increasing concentration of Braco-19. The data recorded at each step was then used to generate stack plots to visualise the changes made in G4 structure upon the addition of Braco-19.

### **Results:**

#### 1. NMR spectra of the RSV-DNA G4s under study:

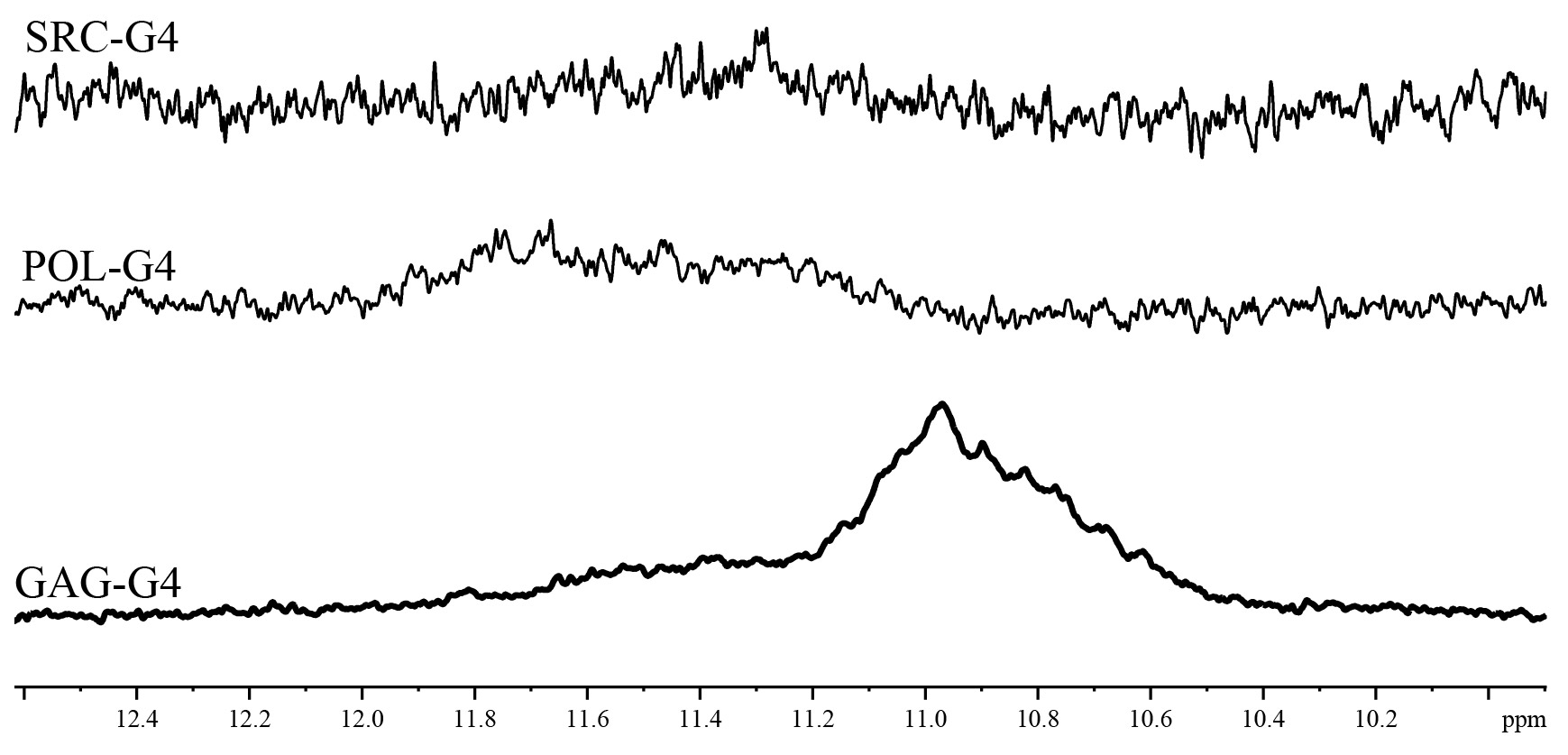

**Supplementary information, Fig 1: 1-D NMR spectra of SG4, PG4, and GG4.** G-quadruplexes are characterised by the formation of Hoogsteen hydrogen bonds between N2-N7 and N1-O6. Formation of G4 structure therefore results in distinct imino proton peaks between 10.0-12.5ppm in NMR spectra. As expected, all sequences under study show peaks in this region, confirming G4 formation. The peaks for SG4 are quite minor as expected from the low G-score of this sequence.

#### 2. Molar CD-spectra of the RSV-DNA G4s:

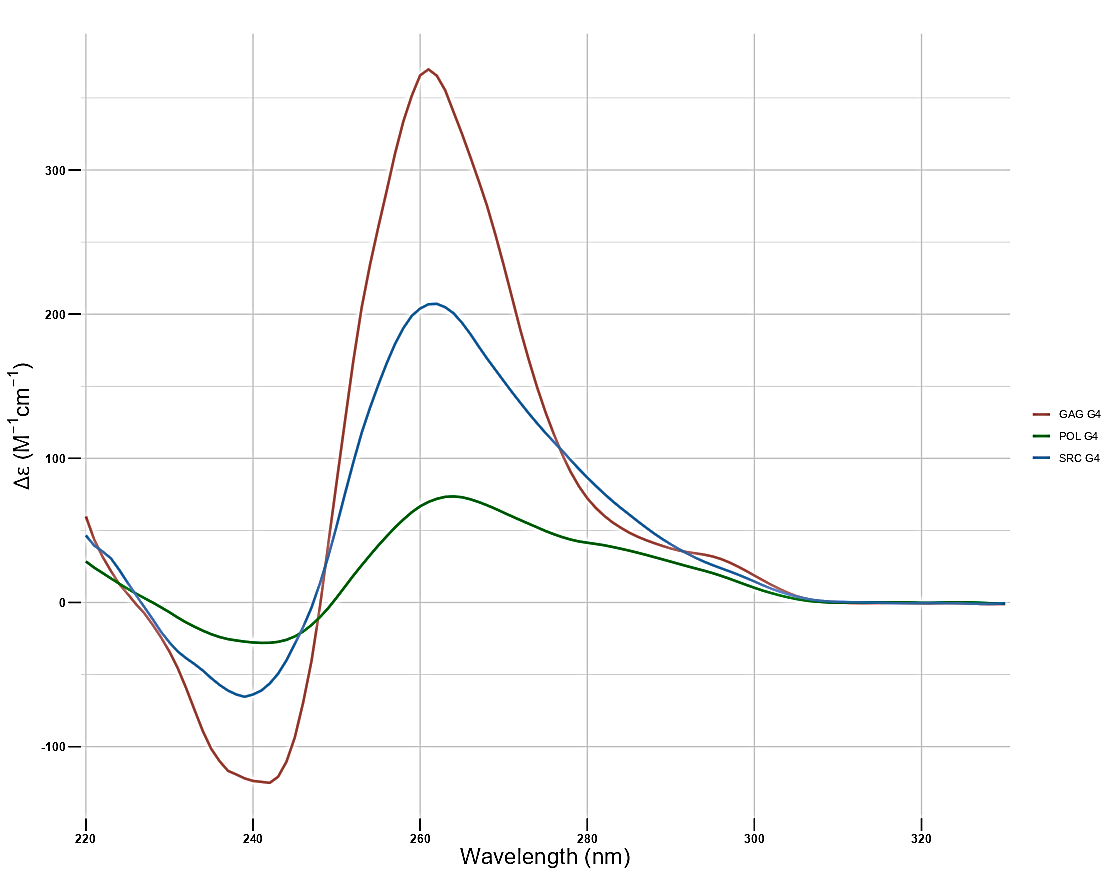

**Supplementary information, Fig 2: Normalized CD-spectra of SG4, PG4 and GG4.** The CD spectra of each sequence was normalized to Δε (M^-1^·cm^-1^) =θ/(32980*c*l), where, Δε is the molar circular dichroism, θ is the ellipticity in millidegrees, c is DNA concentration in mol/L, and l is the path length in cm. The data so obtained was the molar CD spectra of the sequences under study. This spectra was further analysed for the PCA and SVD analysis.

#### 3. Secondary and tertiary structural parameters of the RSV-DNA G4s:

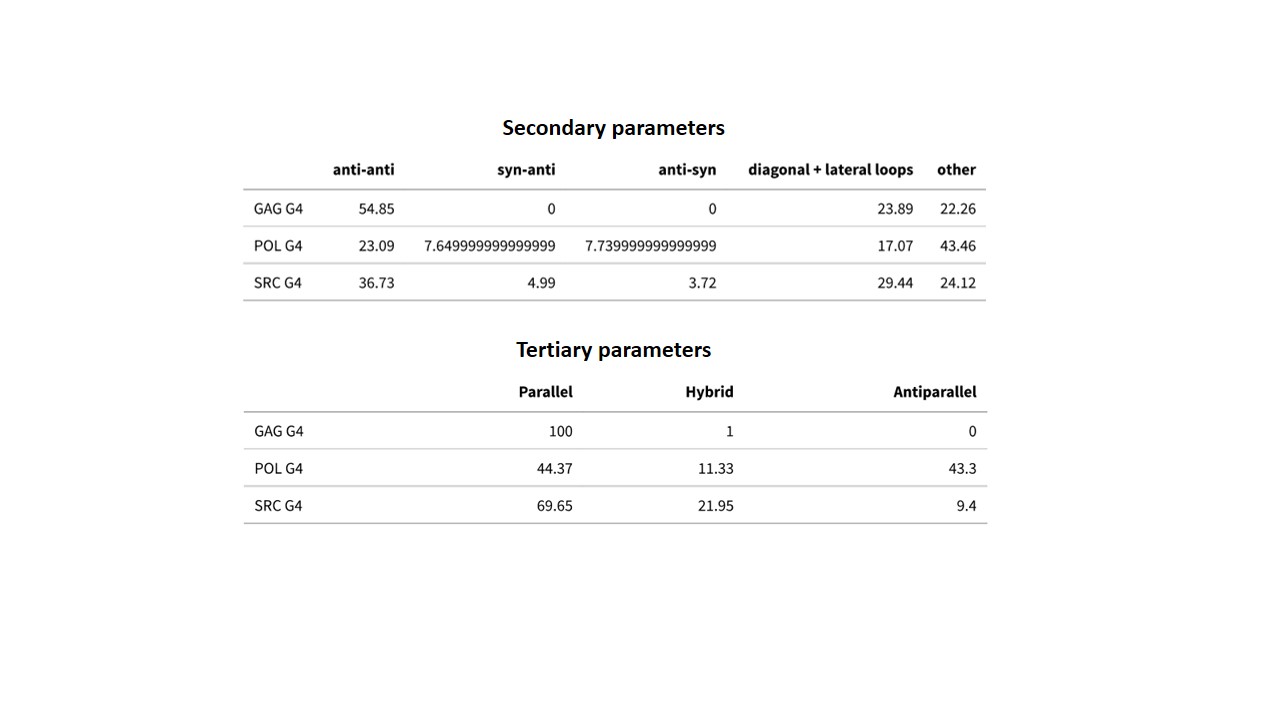

**Supplementary information, Fig 3: Structural parameters of the RSV-DNA G4s.** The normalized CD spectra of SG4, PG4, and GG4 was analysed to obtain the structural topology of these G4s. The CD spectra was analysed against a reference library of 23 G4s using PCA and SVD analysis via R software. GG4 has no syn-anti or anti-syn bonds and is thus completely parallel in nature, while PG4 and SG4 form dynamic G4 structures which remain in equilibrium with each other.

#### 4. DMS Protection assay using wild type sequences and list of mutations:

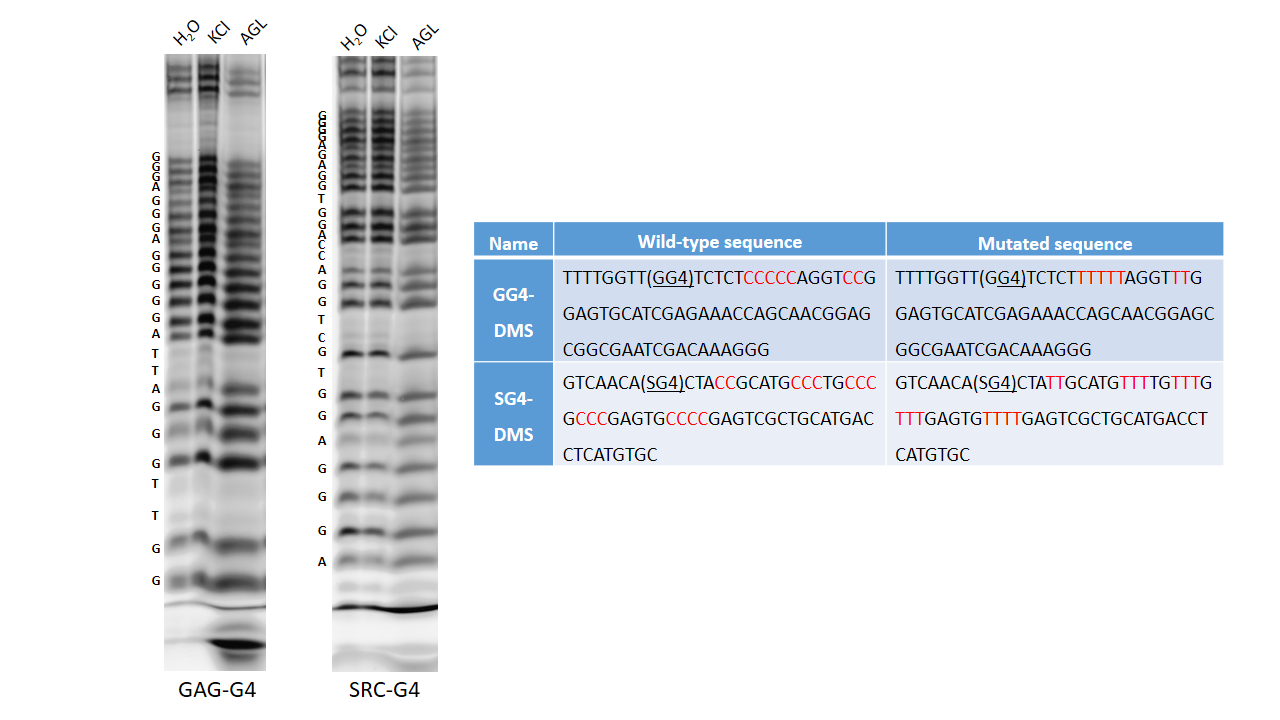

**Supplementary information, Fig 4: DMS Protection assay using GG4 and SG4 in wild-type sequence background.** The wild-type sequence did not show significant protection of guanine residues in the presence of KCl, suggesting the inability to form G4-structures. However, other biophysical data indicated G4 formation by these sequences. Thus, the wild-type sequence following GG4 and SG4 was mutated to reduce cytosine residues which might be involved in hairpin formation.

#### 5. Corroborating ligand binding to RSV-DNA G4s:

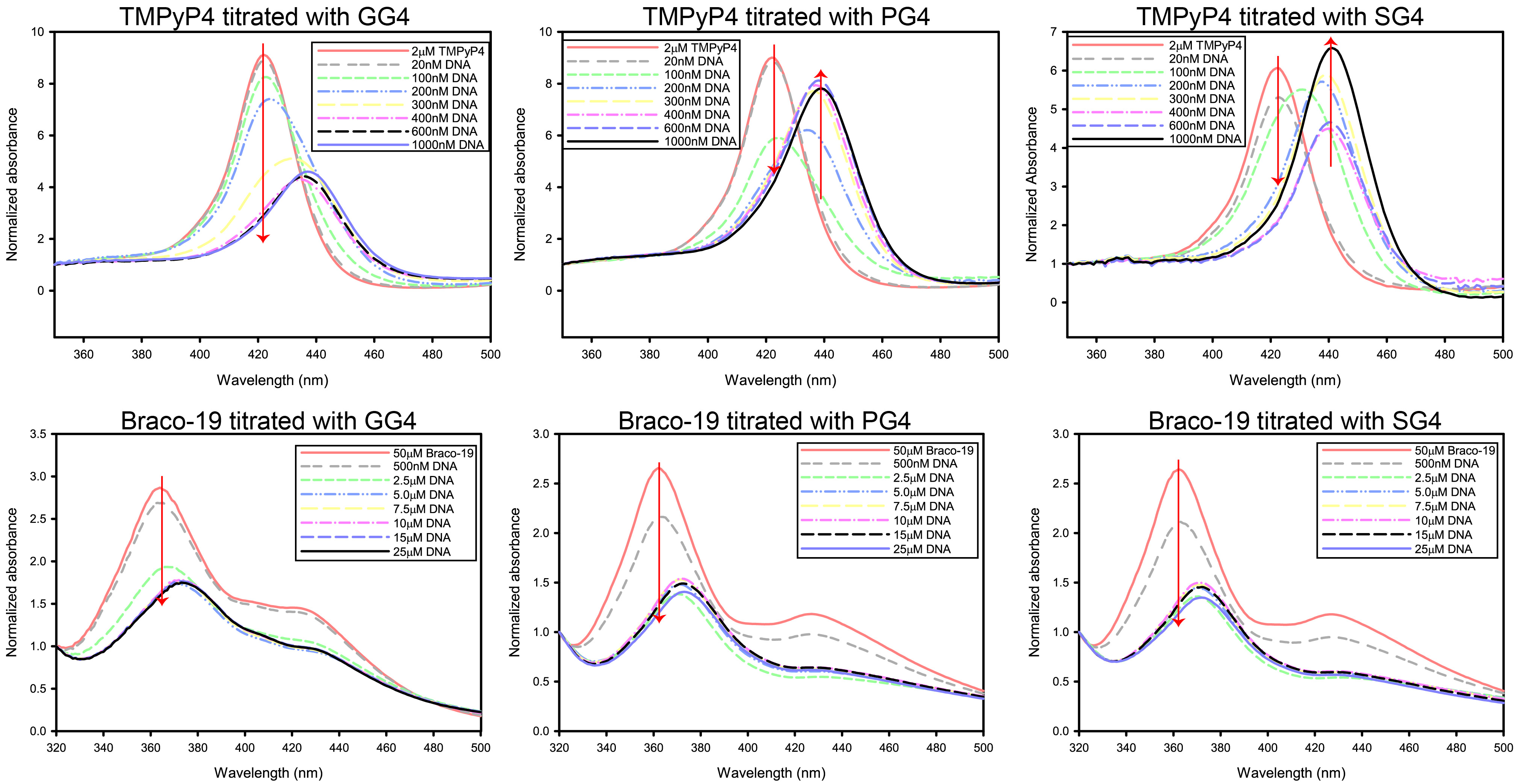

**Supplementary information, Fig 5: Titration of TMPyP4 and Braco-19 with RSV-DNA G4s observed by absorbance spectroscopy.** Both ligands bind the sequences under study as expected, leading to changes in absorbance intensity and maxima.

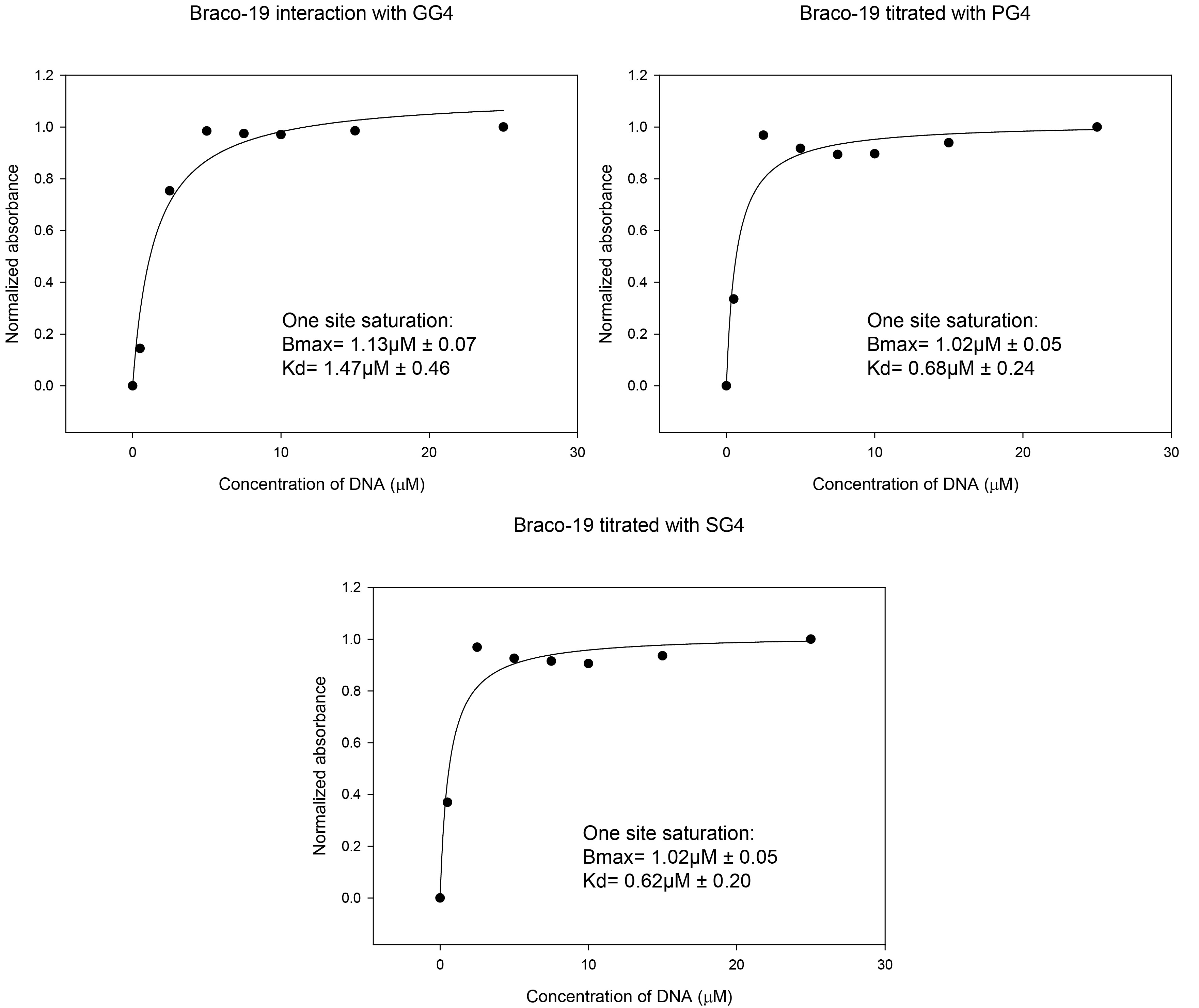

**Supplementary information, Fig 6: Calculation of putative Kd for the interaction between Braco-19 and RSV-DNA G4s.** The change at the absorbance maxima of Braco-19 was noted from the titration experiment and was used to calculate the normalized absorbance. This was plotted as a function of DNA concentration and the graph was fitted using the Sigma Plot software which generated the putative Bmax and Kd of each interaction.

#### 6. Effect of Braco-19 binding on G4 structure:

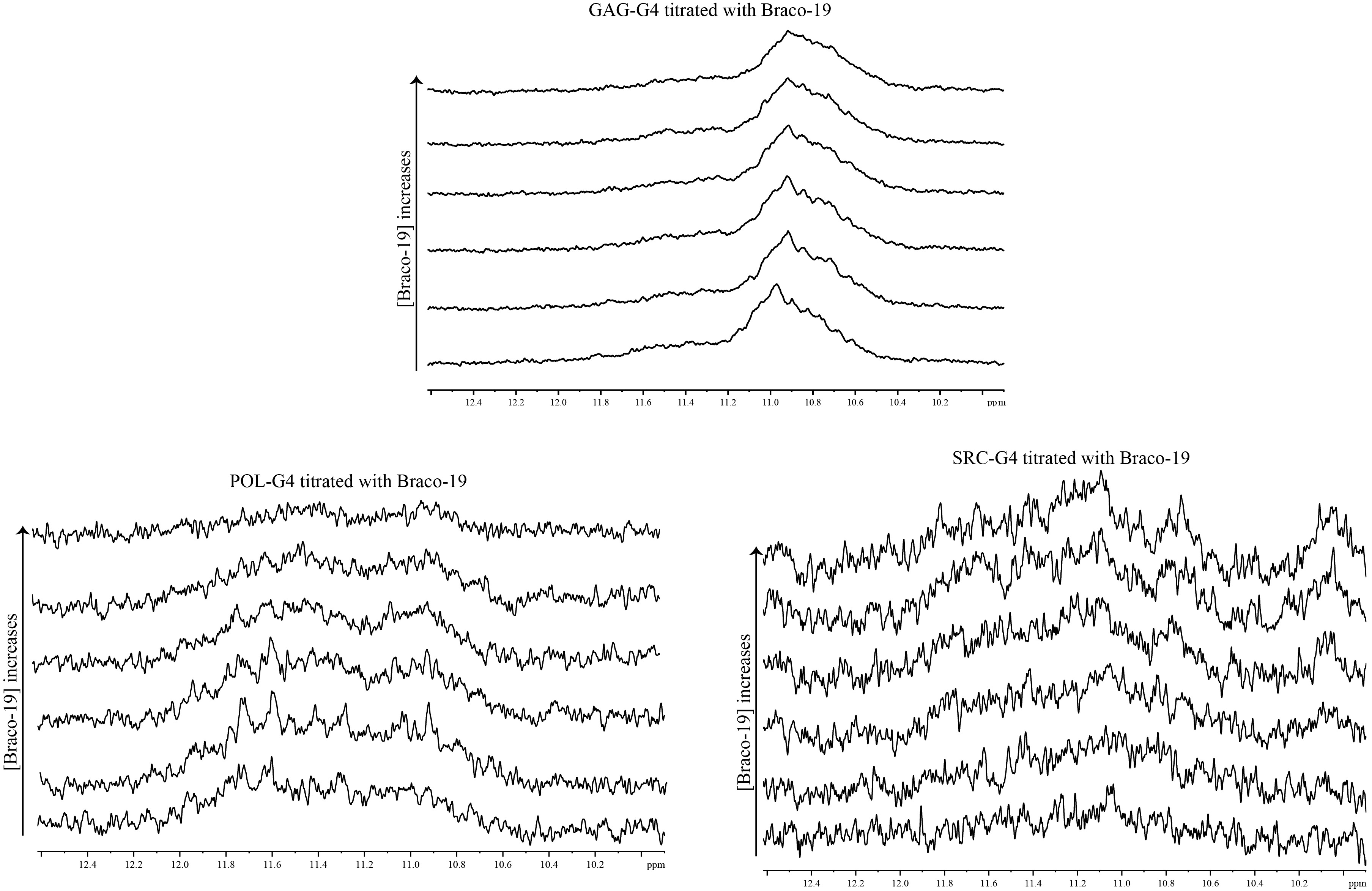

**Supplementary information, Fig 7: Titration of RSV-DNA G4s with Braco-19 induces structural changes.** In the case of GG4, Braco-19 leads to removal of certain peaks suggesting either destabilization or increased dynamicity. However, titration of PG4 or SG4 with Braco-19 led to the development of additional imino proton peaks in the 10-12.5ppm region corresponding to G4 formation. This suggests that these two G4s are stabilized by ligand interaction in addition to undergoing structural alterations. However, this needs to be further corroborated.

#### 7. Homology modelling shows high similarity of chicken nucleolin to human nucleolin

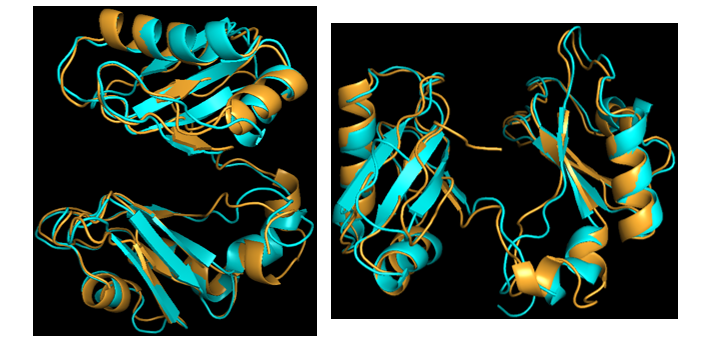

**Supplementary information, Fig 8: Homology model of chicken nucleolin obtained from the I-TASSER software.** Homology modelling of chicken NCL (RBD 1 and 2) with respect to human nucleolin shows high degree of structural similarity. The chicken nucleolin is shown as orange and the human nucleolin is blue.
